# Dark Microglia Are Abundant in Normal Postnatal Development, where they Remodel Synapses via Phagocytosis and Trogocytosis, and Are Dependent on TREM2

**DOI:** 10.1101/2024.10.15.618087

**Authors:** Haley A. Vecchiarelli, Kanchan Bisht, Kaushik Sharma, Sammy Weiser Novak, Marianela E. Traetta, Monica Emili Garcia-Segura, Marie-Kim St-Pierre, Julie C. Savage, Cory Willis, Katherine Picard, Maude Bordeleau, Nathalie Vernoux, Mohammadparsa Khakpour, Rishika Garg, Sophia M. Loewen, Colin J. Murray, Yelena Y. Grinberg, Joel Faustino, Torin Halvorson, Victor Lau, Stefano Pluchino, Zinaida S. Vexler, Monica J. Carson, Uri Manor, Luca Peruzzotti-Jametti, Marie-Ève Tremblay

**Affiliations:** Division of Medical Sciences, University of Victoria, Victoria, British Columbia, Canada; Axe neurosciences, Centre de recherche du CHU de Québec-Université Laval, Québec, QC, Canada; Waitt Advanced Biophotonics Core, Salk Institute for Biological Studies, La Jolla, CA, USA; Neuroscience Graduate Program, University of Victoria, Victoria, British Columbia, Canada; Department of Clinical Neurosciences and NIHR Biomedical Research Centre, University of Cambridge, Cambridge, UK; Department of Metabolism, Digestion and Reproduction, Imperial College London; United Kingdom; Westview High School, San Diego, CA, USA; Division of Biomedical Sciences, University of California – Riverside, Riverside, CA, United States; Department of Neurology, University of California – San Francisco, San Francisco, CA, United States; Department of Cell & Developmental Biology, School of Biological Sciences, UC San Diego, La Jolla, CA; Department of Molecular Medicine, Université Laval, Québec City, Québec, Canada; Neurology and Neurosurgery Department, McGill University, Montréal, Québec, Canada; Department of Biochemistry and Molecular Biology, The University of British Columbia, Vancouver, British Columbia, Canada; Centre for Advanced Materials and Related Technology (CAMTEC), University of Victoria, Victoria, BC, Canada; Institute for Aging and Lifelong Health (IALH), University of Victoria, Victoria, BC, Canada

## Abstract

This study examined dark microglia—a state linked to central nervous system pathology and neurodegeneration—during postnatal development in the mouse ventral hippocampus, finding that dark microglia interact with blood vessels and synapses and perform trogocytosis of pre-synaptic axon terminals. Furthermore, we found that dark microglia in development notably expressed C-type lectin domain family 7 member A (CLEC7a), lipoprotein lipase (LPL) and triggering receptor expressed on myeloid cells 2 (TREM2) and required TREM2, differently from other microglia, suggesting a link between their role in remodeling during development and central nervous system pathology. Together, these results point towards a previously under-appreciated role for dark microglia in synaptic pruning and plasticity during normal postnatal development.

## Main Text

Crucial for the proper development, function and plasticity of the central nervous system (CNS) are microglia, the CNS’s resident innate immune cells^1^. Microglia contribute to the formation and maturation of dendritic spines and the elimination of synapses by stripping (i.e., physical separation of pre- and post-synaptic elements by their intervening processes)^2,3^ or pruning (i.e., direct synaptic engulfment). Pruning, which involves phagocytosis or trogocytosis (nibbling) of synaptic structures^4,5^, plays an important role in learning, memory, and sociability, among other behaviors^1,6,7^. Indeed, aberrant levels of microglial synaptic modulation during early postnatal development have been linked to neurodevelopment conditions^7,8^.

Microglia are an intrinsically heterogenous population of cells, which exert diverse functions in health and disease by acquiring distinct states^9,10^. Recently, we identified an unique microglial state, only detectable by electron microscopy (EM)^11^, dark microglia (DM). DM differ from non-dark, typical microglia by their loss of the microglial nuclear heterochromatin motif, as well as reduced levels of the canonical microglia markers ionized calcium-binding adapter molecule 1 (IBA1) and CX3C motif chemokine receptor 1 (CX3CR1)^11^; DM display additionally ultrastructural markers of cellular stress including a dark/electron-dense appearance in EM, but also dilation of the endoplasmic reticulum and Golgi apparatus, and mitochondrial cristae alteration^11^. Notably, while DM are rare in healthy young adult mice, they can increase in number up to 10-fold upon exposure to various challenges, including chronic psychological stress^11^, aging^11^, fractalkine signaling deficiency^11^, maternal immune activation^12^, and Alzheimer’s^13,14^ and Huntington’s^15^ disease pathologies. However, the relevance of DM during normal physiological conditions is still unknown. Herein, we discovered and investigated the occurrence of DM in C57BL/6J mice during normal brain development (postnatal days (P)5-15) via a combination of scanning and transmission electron microscopy (SEM and TEM), volumetric focused ion beam (FIB)-SEM, and imaging mass cytometry (IMC).

EM analysis revealed an abundance of DM in the ventral hippocampus (vHIP) *cornu ammonis (CA)1 stratum radiatum and stratum lacunosum moleculare*, a known hot spot for these cells^11^, during the first two weeks of postnatal life (**Fig.1A-B**). DM showed a distinctive electron-dense cytoplasm and nucleoplasm and loss of microglia-distinctive nuclear heterochromatin, as previously observed in DM during pathology^11^ (**Fig.1A-D**). DM in development were observed in contact with the neuropil, neurons, astrocytes, typical microglia (**Fig.1E**) and blood vessels (**Fig.1B**), and exhibiting satellite relationships juxtaposing neurons (**Fig.1C**)—a hallmark of microglial-mediated regulation of neuronal activity^16^.

**Figure 1.**
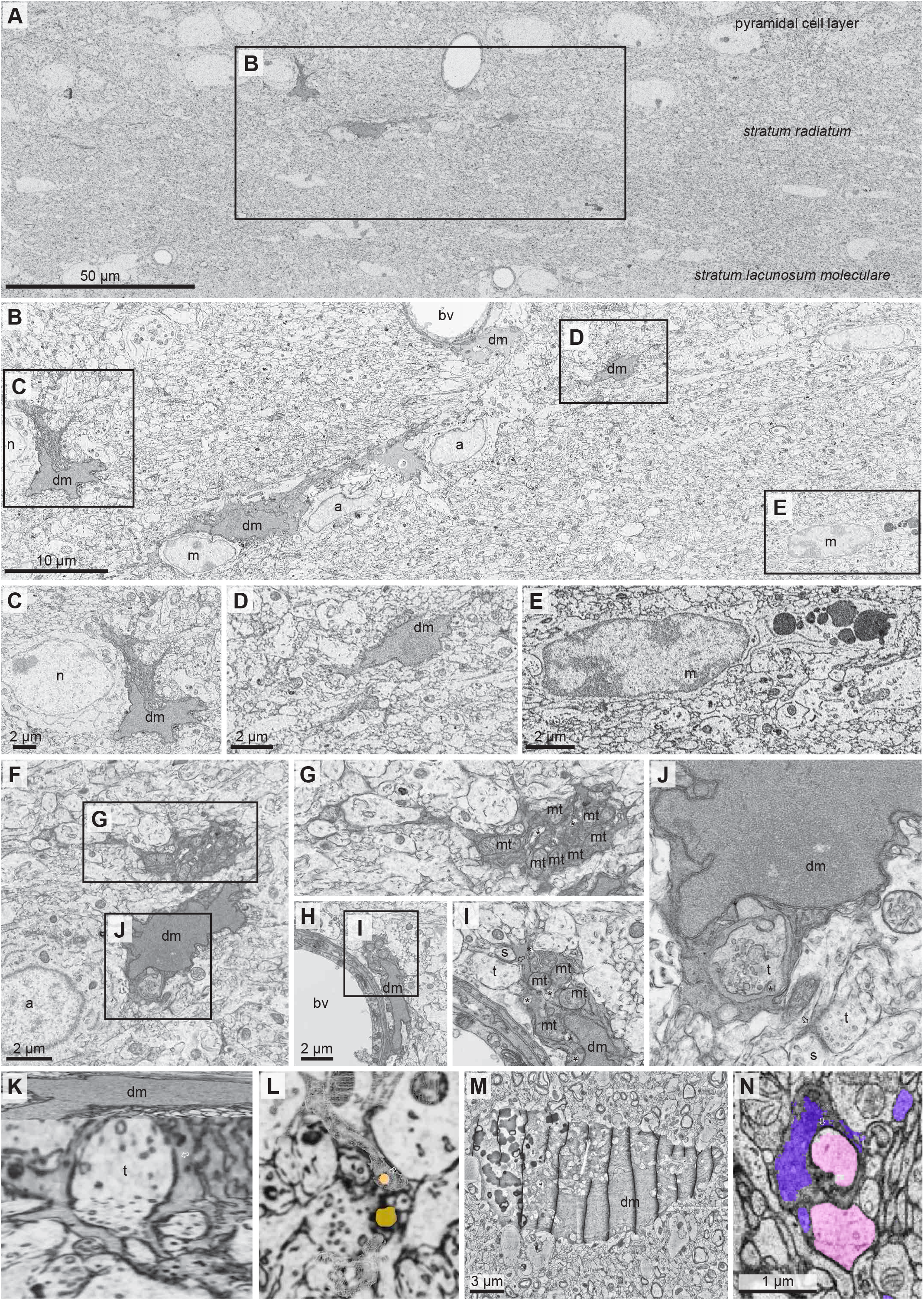
Dark microglia are present in the ventral hippocampus CA1 region during normal early postnatal development, interacting with synapses and the vasculature, performing phagocytosis and trogocytosis. A: Scanning electron microscopy imaging of the ventral hippocampus CA1 of P10 C57BL/6J mice showing the distribution of dark microglia (dm) and typical microglia (m) across *strata pyramidale, radiatum* and *lacunosum-moleculare*. B: Higher magnification view of four dark microglia (one of which contacts an interneuron (n), one of which directly associates with a blood vessel (bv) and one of which juxtaposes one typical microglia and two astrocytes). C-E: Higher magnification views of two dark and one typical microglia, respectively. C: The dark microglia observed here occupies a satellite position to an interneuron, directly touching its cell body. F-I: Transmission electron microscopy revealed dark microglia present cellular stress markers, including endoplasmic reticulum dilation and accumulation of mitochondria, sometimes with altered cristae structure. F: Dark microglia juxtaposing an astrocytic cell body. G: Higher magnification view of a proximal process showing an accumulation of mitochondria (mt) and endoplasmic reticulum dilation (asterisks). H: Dark microglia located next to a blood vessel, directly apposing the basement membrane. I: Higher magnification view of the dark microglia showing numerous mitochondria and endoplasmic reticulum dilation (asterisks). In addition, a dark microglial ramified process interacts with a synapse, contacting a dendritic spine (s) and an axon terminal (t) (arrowhead). J: Higher magnification view of (F) showing the synaptic vesicles in the axon terminal (t) and dilated endoplasmic reticulum (asterisk) in the dark microglia cell body. A distal process from the same dark microglia is found in contact with a synapse (arrow), touching a dendritic spine (s) and an axon terminal (t). K-L: Stills of a dark microglia (dm) imaged by focused-ion beam-scanning electron microscopy in the ventral hippocampus CA1 of P10 C57BL/6J mice from Supplemental Video 1. K: DM shown in 3D along xyz planes, phagocytosing an axonal terminal (t). L: Dark microglia process (outlined), contacting a presynaptic axon terminal with a corresponding dendritic spine (yellow) with a post-synaptic density. This dark microglial process is performing trogocytosis (nibbling) at an axonal terminal (orange). M: Example of a dark microglia in cortical layer V from the anterior temporal cortex from the H01 data set^17^ in Neuroglancer (segmentation IDs: 40881443277 and 85062801060). N: dark microglial process (purple) interacting with an axon segment (segmentation ID: 5013063676; pink), potentially engulfing a portion of it.Representative electron micrographs. Scale bars are indicated on the electron micrographs. a=astrocyte; bv=blood vessel; dm=dark microglia; m=microglia; mt=mitochondria; n=neuron; s=dendritic spine; t=axon terminal. * asterisks indicate examples of endoplasmic reticulum dilation; arrow indicates dark microglial process in contact with synapses.

We next investigated additional features of DM in the postnatal vHIP, finding that they also displayed endoplasmic reticulum dilation and accumulation of mitochondria with altered cristae accumulation (**Fig.1F-I**). Additionally, DM processes made extensive contacts with synapses (**Fig.1F-J**), another hallmark of DM in pathology, and exhibited various types of intracellular inclusions, including synaptic elements (**Fig.1J-K**). FIB-SEM with an isotropic resolution of 14 nm, was utilized to study in 3D the intracellular synaptic inclusions in DM from the CA1 *stratum lacunosum-moleculare* at P10 (**Video.1**). This cutting-edge technique revealed different modalities of interactions between DM and synaptic elements, with DM both engulfing axonal terminals completely (**Fig.1K** and **Video.1**) or nibbling small pieces of axon terminals via trogocytosis (**Fig.1L** and **Video.1**). This novel function of DM was further explored in a publicly available dataset (H01) of a reconstructed adult human temporal cortex^17^, in which DM were detected (**Fig.1M**), and showed processes interacting with, and potentially engulfing, an axonal segment of a pyramidal neuron (**Fig.1M-N**). This intriguing finding demonstrates that DM are active participants in synaptic remodeling, engaging in both phagocytosis and trogocytosis, in mice during development, but also perform similar synaptic interactions in humans, although more work across the developmental trajectory is necessary.

We followed up our ultrastructural data by utilizing a targeted approach for assessing phenotypic markers of DM in development. Immunoreactivity against the homeostatic microglial marker transmembrane protein 119 (TMEM119)^18^ was not detected on DM in development, contrary to strong signal observed in neighboring typical microglia (**Fig.2A-B**) and DM in development were strongly immunopositive for cluster of differentiation molecule 11b (CD11b) (**Fig.2C-D**), an essential component of complement receptor 3 important for synaptic pruning^19^, as was observed for DM in pathology. CD11b was observed in DM at the plasma membrane of their distal processes encircling synaptic elements (**Fig.2C-D**). Newly, we observed that these DM in development are immunoreactive for C-type lectin domain family 7 member A (CLEC7a)/Dectin-1 (**Fig.E-F**), an anti-fungal pattern recognition receptor, and for lipoprotein lipase (LPL) (**Fig.2G-H**), an enzyme involved in phagocytosis and metabolism of lipoproteins^20^.

**Figure 2.**
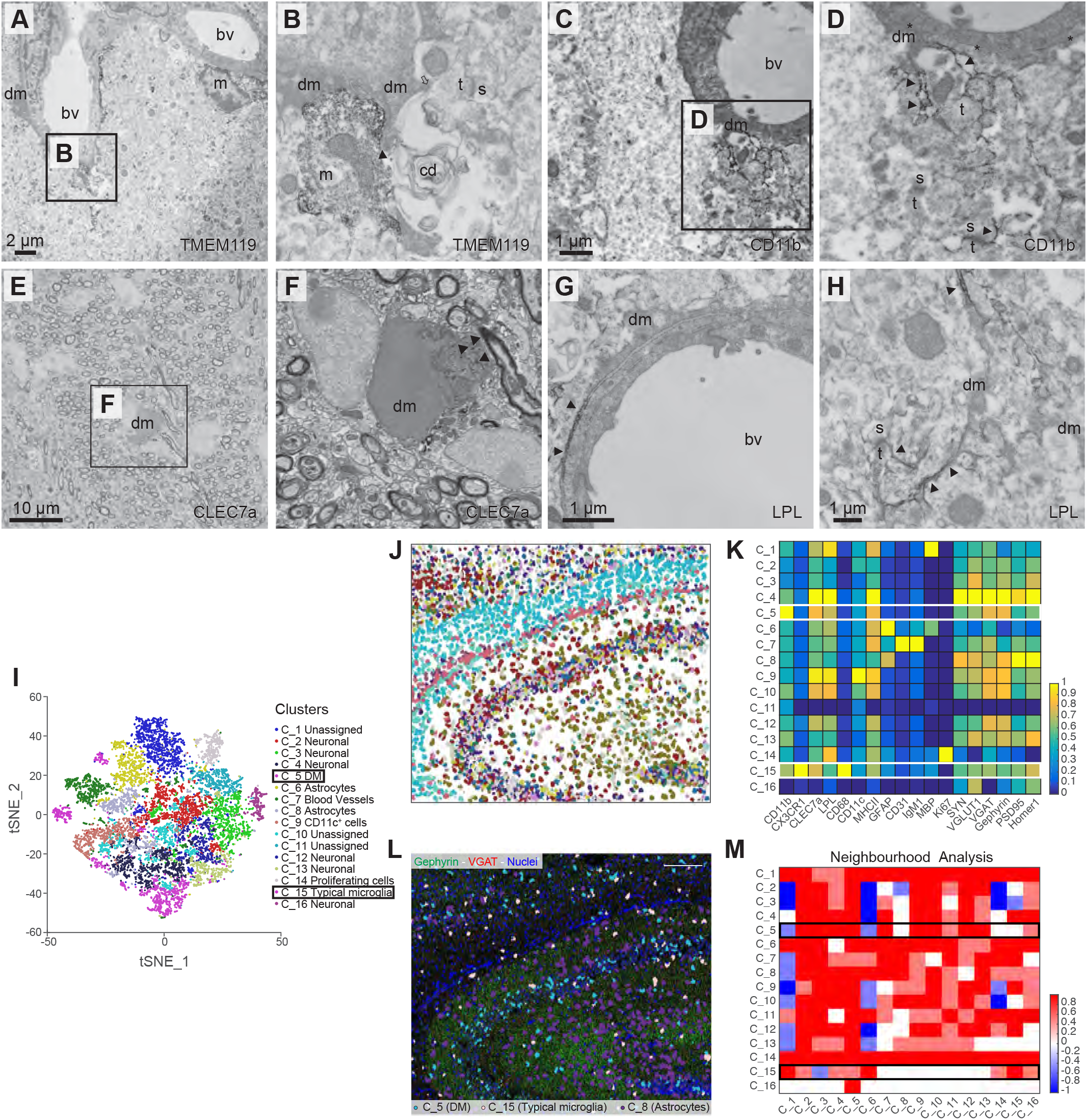
Dark microglia in development do not express the typical microglia marker TMEM119, yet express CD11b, CLEC7a and LPL. A-B: Using immunocytochemical transmission electron microscopy, dark microglia in development in the ventral hippocampus CA1 *strata radiatum* and *lacunosum-moleculare* of P10 C57BL/6J mice were not found to express TMEM119. A: Dark microglia (dm) process ensheathing a blood vessel (bv). A typical microglia (m) cell body stained for TMEM119 is also seen contacting a nearby blood vessel. B: Higher magnification view of the dark microglia process showing direct juxtaposition (arrowhead) with a typical microglia process stained for TMEM119. This dark microglia process is also touching (arrow) an axon terminal (t) making a synapse onto a dendritic spine (s). Both the dark and typical microglia processes are additionally associated with an extracellular space pocket containing partially degraded cellular elements or debris (cd). C-D: Example of dark microglia stained for CD11b observed in the ventral hippocampus CA1 *strata radiatum* and *lacunosum-moleculare* of P10 C57BL/6J mice using transmission electron microscopy. C: The dark microglia process is seen surrounding a blood vessel and juxtaposing a dendrite (d). D: Higher magnification view of the process showing CD11b staining on its distal extremities (arrowheads), where touching or encircling neuronal elements, such as dendritic spines (s) and axon terminals (t). E-F: Example of dark microglia stained for CLEC7a obtained using scanning electron microscopy in P15 C57BL/6J mice. F: Higher magnification view of the dark microglia with immunoreactivity for CLEC7a (arrowheads). G-H: Examples of dark microglia stained for LPL observed in the ventral hippocampus CA1 of P10 C57BL/6J mice using transmission electron microscopy. G: A dark microglia wrapped around a blood vessel display positive immunostaining for LPL in its processes (arrowhead). H: Dark microglial processes immunopositive for LPL (arrowheads) near a dendritic spine (s) and axon terminal (t).Representative electron micrographs. Scale bars are indicated on the electron micrographs. bv=blood vessel; cd=cellular debris; dm=dark microglia; m=microglia; s=dendritic spine; t=axon terminal. I-M: Imaging mass cytometry (IMC) I: Unsupervised cluster (C_1-16) Phenographs obtained from IMC analysis of hippocampus *cornu ammonis* (CA)1 at P14, visualised on a t-SNE plot. J: Representative picture of the different spatial distribution of C_1-16 via pseudocolouring of single hippocampal cells. K: Heatmap showing the relative mean expression of the 18 markers used in the IMC analysis. Marker attribution identified putative cell types for the following clusters: dark microglia (C_5), typical microglia (C_15), MBP^+^ myelin structures (C_1), neuronal clusters (C_4,12,13), CD31^+^IgM+ blood vessels (C_7), GFAP^+^ astrocytes (C_6,8), CD11c^+^ dendritic cells (C_9), KI67^+^ proliferating cells (C_14). Some clusters (C_2,3,10,11,16) could not be unequivocally assigned to a specific cell type. L: Representative picture showing different spatial distribution of the C_5 dark microglia, C_15 typical microglia and C_8 astrocytes in relation to gephyrin (green) and VGAT (red) expression. Nuclei are in blue. M: Heatmap displaying spatial interactions between Phenograph-derived clusters as revealed by neighbourhood analysis. Rows 5 and 15 are boxed, showing the cell clusters (columns) in the neighbourhood of C_5 (dark microglia) and C_15 (typical microglia). Colors indicate the prevalence of cell-type interactions across the region of interest, with blue squares representing avoidance and red representing positive interactions.

We then expanded on these data using an IMC approach, which allowed us to simultaneously use 18 different antibodies on the P14 vHIP to obtain single cell spatial data (**Fig.2I-M**). Over the total of 16 clusters (C_1-16) that were identified, we found a cluster of CD11b^high^/CX3CR1^low^ cells (C_5), which was distinct from CX3CR1^high^ typical microglia (C_15) and represented a putative DM cluster (**Fig.2K**). Compared to C_15 typical microglia, C_5 DM also expressed higher levels of MHCII, as shown for DM in pathology^11^, and CLEC7a and LPL, which we newly found on DM in development (**Fig.2E-H**).

IMC also allowed us to test the expression of non-microglial proteins in DM and identify the spatial relationship between DM and other cell clusters. We found that C_5 DM showed increased co-expression of the pre- and post-synaptic markers VGAT and gephyrin **(Fig.2K-L)**, structural components of inhibitory synapses. Complementary unsupervised neighbourhood analysis revealed that C_5 DM were strongly interacting with CD31^+^/IgM^+^ blood vessels (C_7), which was not observed for C_15 typical microglia **(Fig.2M)**. We also found that neuronal clusters defined by the expression of axonal and dendritic markers (C_2,3,4,12,13, 16) predominantly localized to the vicinity of C_5 DM. In contrast, interactions between neuronal clusters and C_15 typical microglia were reduced (C_2,3,4,16) or absent (C_12,13). These data, combined with our FIB-SEM evidence of phagocytosis and trogocytosis, further support a putative functional role of DMs’ interactions with neurons, synaptic elements and the vasculature in the early postnatal vHIP.

This phenotypic characterization reveals that DM in development display a similarity to other microglial states, such proliferation-associated microglia (PAM)^21^, disease-associated microglia (DAM)^22^ and microglia with a neurodegenerative phenotype (MgND)^23^. As such, we next investigated the expression of triggering receptor expressed on myeloid cells 2 (TREM2) on DM in development, especially given the requirement of TREM2 for the emergence of different states of microglia microglia^20^—including DAM^22^ and MgND^23^—and, that TREM2 regulates LPL expression as part of a broader shift in microglial metabolism and function^22^. DM in development are immunoreactive for TREM2 (**Fig. 3A-B**). Upon analysing TREM2 knock-out (KO) mice (**Fig.3C**), we found a near total loss of DM at P15 compared to wild-type mice (**Fig.3D**), with no change in typical microglial density (**Fig.3E**).

**Figure 3.**
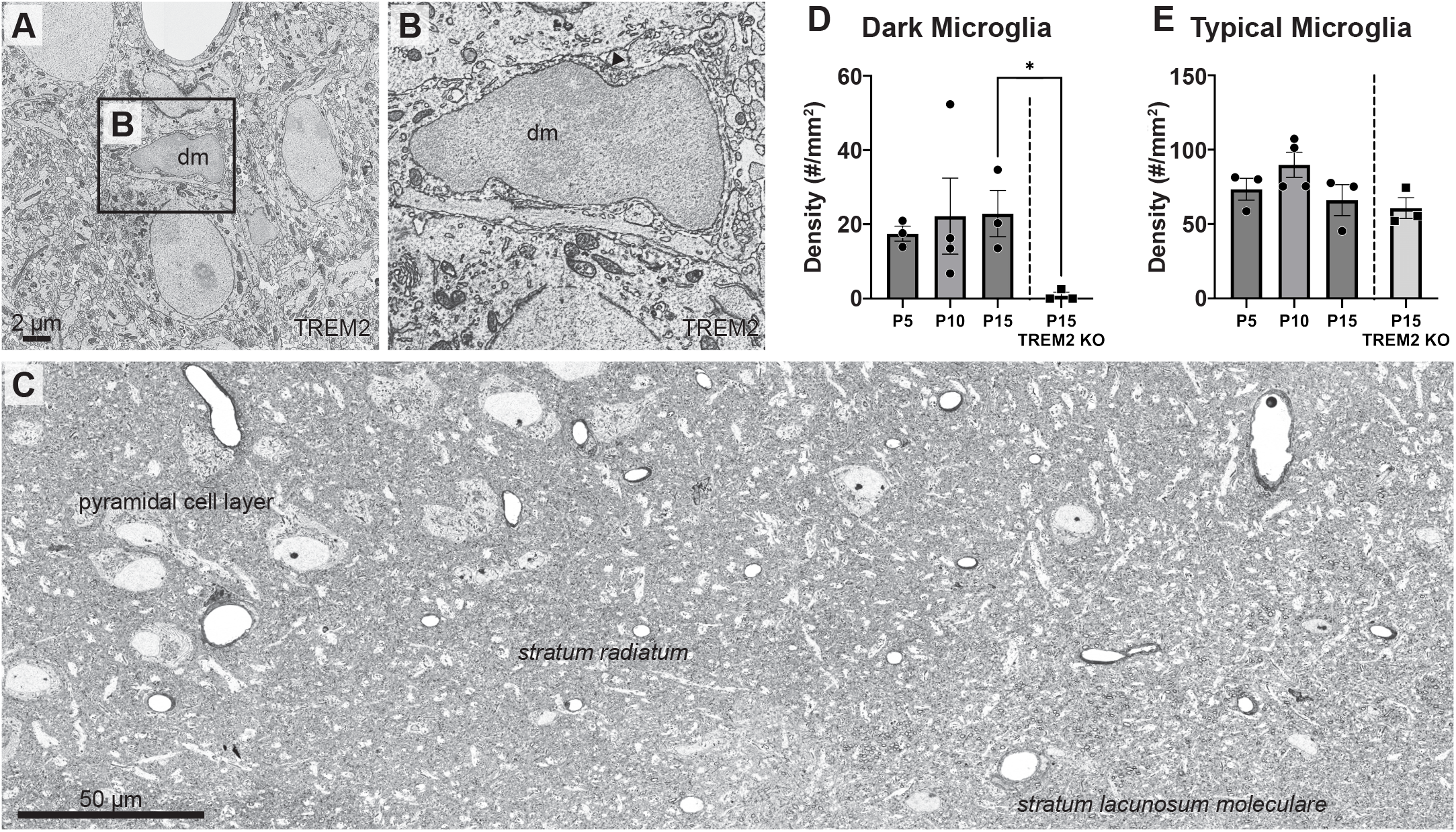
Dark microglia in development are dependent on TREM2. A-B: Example of an intermediate dark microglia from P10 C57BL/6J mice showing TREM2 immunopositivity in the ventral hippocampus dentate gyrus using scanning electron microscopy. A: A dark microglia cell in the ventral hippocampus dentate gyrus. B: Higher magnification view of (B) the intermediate dark microglia cell body showing TREM2 staining (arrowhead). C: Representative scanning electron microscopy micrograph from P15 TREM2 KO mice in the ventral hippocampus CA1 region containing the *strata pyramidale, radiatum* and *lacunosum-moleculare*. No dark microglia are present.Representative electron micrographs. Scale bars are indicated on the electron micrographs.dm=dark microglia; t=axon terminal; s=dendritic spine. arrow indicates dark microglial process in contact with synaptic elements. D: Quantification of dark microglial density across the ventral hippocampus CA1 combined *stratum radiatum* and stratum *lacunosum-moleculare*, with no statistically significant differences between P5, P10 and P15 (F(2,7)=0.12; p=0.89; R^2^=0.03). There was a significant reduction of dark microglia in TREM2 KO mice, compared to wild type at P15 (t(4)=3.50; p=0.03; R^2^=0.75). E: Quantification of typical microglial density across the ventral hippocampus CA1 combined *stratum radiatum* and stratum *lacunosum-moleculare*, with no significant differences between P5, P10 and P15 (F(2,7)=2.01; p=0.20; R^2^=0.36). There were no significant differences between the number of typical microglia in TREM2 KO at P15 compared to wild type mice (t(4)=0.43; p=0.69; R^2^=0.04).Statistical analysis for comparison between P5, P10 and P15 was performed using a one-way ANOVA, and F-, p-and R^2^-values were reported. Statistical analysis for comparison between P15 wild type and TREM2 KO were performed using a two-tailed Student’s t-test, and t-, p- and R^2^-values were reported. n=3-4 animals/group. *p<0.05.

Altogether this work reveals that DM are abundant during early postnatal development at a time when gliogenesis, synaptic density and synaptic pruning peak^24^. Here, DM display similar ultrastructural cellular stress markers, association with the vasculature and neurons (including synaptic elements) as DM found in pathology. Using cutting-edge volumetric FIB-SEM, we observed for the first-time phagocytosis and trogocytosis of axonal terminals by a DM. Additionally, we showed that DM in development possess an immunophenotype similar to other altered microglial states (i.e., PAM, DAM and MgND), including CLEC7a, LPL and TREM2 positivity, and that they are dependent on TREM2 expression.

The abundance of DM during developmental sensitive periods, their localization, phenotypic expression and observed functioning (e.g., phagocytosis and trogocytosis) suggests that DM in development are involved in essential physiological processes such as vascular and synaptic remodeling. In conclusion, while the abundance of DM during development point to their role in critical processes like synaptic remodeling, further research is necessary to elucidate the precise role of DM phagocytosis and trogocytosis, and the importance of DM in development in both healthy and diseased states.

## Materials and Methods

### Animals

Experiments were conducted under the approval of the Université Laval, University of Victoria, University of California San Francisco, University of California Riverside and Cambridge University Institutional Animal Care and Use Committees according to guidelines outlined by the U.S. Department of Agriculture’s Animal Welfare Act and Animal Welfare Regulation, the Canadian Council on Animal Care and the Animals (Scientific Procedures) Act 1986 Amendment Regulations 2012 in the United Kingdom.

Both male and female mice were utilized in all studies from postnatal day (P)5-P15. All experiments except the triggering receptor expressed on myeloid cells 2 (*Trem2^−/−^*) knockout (KO) studies utilized C57BL/6J mice (either from commercial sources (The Jackson Laboratory) or bred in-house from animals originally acquired from commercial sources). Mice were housed under a 12-hour light-dark cycle at 22-25 °C with free access to food and water. For TREM2 and CLEC7a staining, pregnant dams were exposed to interperitoneally to saline or polyinosinic:polycytidylic acid (poly I:C) (5 mg/kg in saline), respectively, at embryonic day (E)9.5 and their offspring utilized for analysis.

### Tissue preparation

For all studies except imaging mass cytometry (IMC), mice were anaesthetized with ketamine/xylazine (80 mg/kg, i.p.) and transcardially perfused with 0.1 % glutaraldehyde in 4 % paraformaldehyde (PFA)^25^ or 3.5% acrolein followed by 4% PFA^26^. 50 μm thick transverse coronal sections of the brain were cut in sodium phosphate buffer (PBS; 50 mM, pH 7.4) using a vibratome (Leica VT1000S) and stored at -20°C in cryoprotectant (ethylene glycol:glycerine:PBS 30:30:40 vol:vol:vol) solution^26^ until further processing. For IMC mice were anaesthetized with ketamine/xylazine (80 mg/kg, i.p.), brains extracted and flash frozen on dry ice. Brain blocks were cryo-sectioned with a 6 μm thickness using a cryostat (OTF5000, Bright Instruments, Huntingdon, United Kingdom) with a microtome blade (A35, Feather, Osaka, Japan). Upon collection on superfrost slides (J1800AMNZ, Epredia Netherlands B.V., Breda, Netherlands), sections were stored at -80 °C until use.

### Immunoperoxidase staining

Immunostaining with specific antibodies against transmembrane protein 119 (TMEM119) (abcam, #ab209064), cluster of differentiation molecule 11b (CD11b) (AbD Serotec, #MCA711GT), C-type lectin domain family 7 member A (CLEC7a) (Invivogen, mabg-mdect), lipoprotein lipase (LPL) (abcam, #ab21356) and triggering receptor expressed on myeloid cells 2 (TREM2) (R&D Systems, #AF1729), was performed as described in **Table 1**. Briefly, the samples were washed with PBS, quenched, blocked and incubated with primary antibodies, washed with PBS, incubated with secondary antibodies and washed. Following secondary antibody staining, sections were incubated with avidin/biotin-based peroxidase complexes (ABC) (1:100 in Tris-buffered saline (TBS); Vector Laboratories, #PK-6100) or streptavidin-horseradish peroxidase (HRP) (1:1000 in blocking buffer; Jackson, #016-030-084) to amplify target antigen signal. The brain sections were afterwards treated with diaminobenzidine (DAB; 0.05%) and hydrogen peroxide (0.015%) to reveal the immunostaining.

**Table 1.**
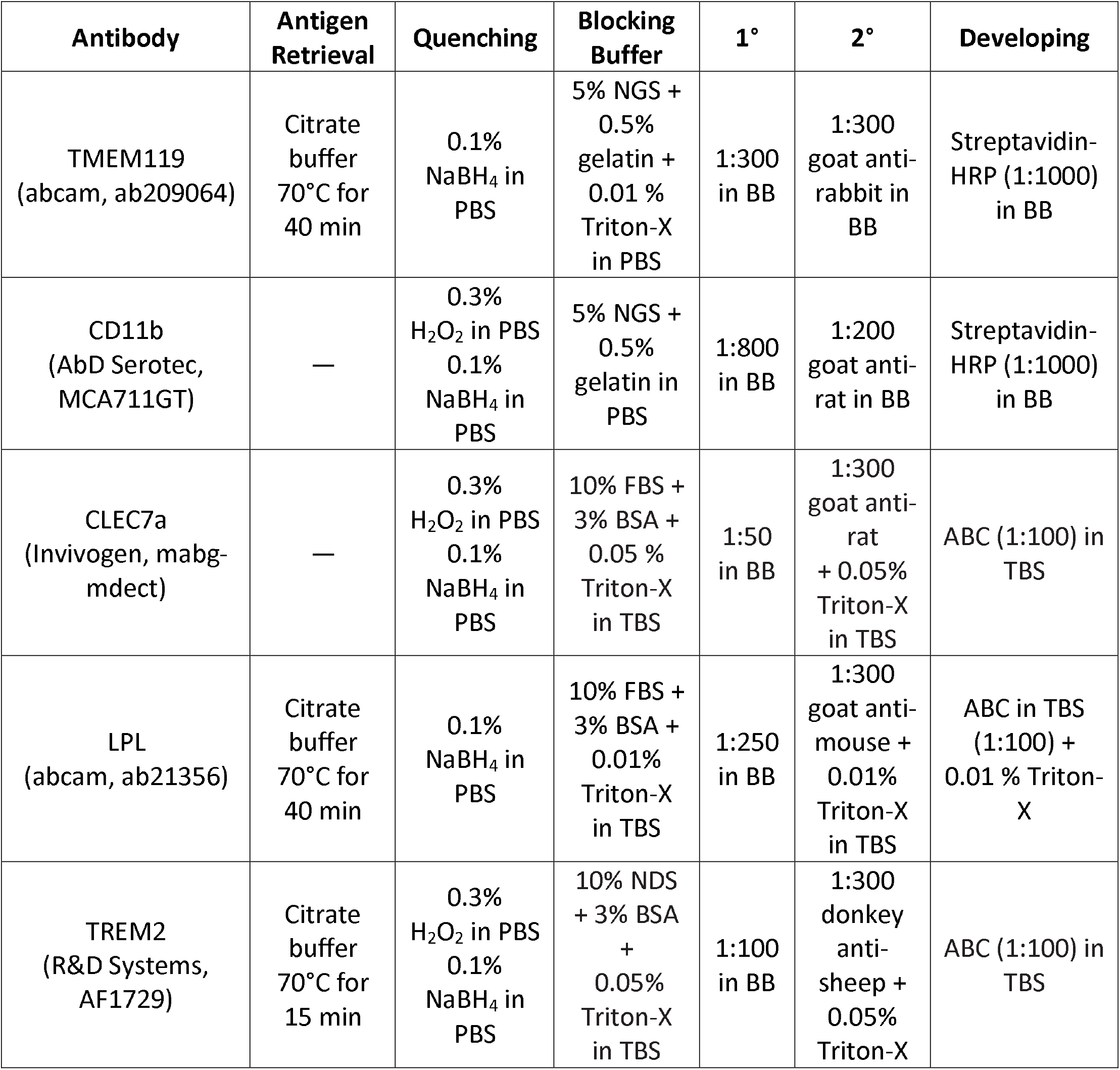
Antibody Conditions. ABC=avidin biotin complex; BB=blocking buffer; BSA=bovine serum albumin; CD11b=cluster of differentiation 11b; CLEC7a=C-type lectin domain family 7 member A; FBS=fetal bovine serum; HRP=horse radish peroxidase; LPL=lipoprotein lipase; NDS=normal donkey serum; NGS=normal goat serum; PBS=phosphate buffered saline; TBS=Tris buffered saline; TMEM119=transmembrane protein 119; TREM2=triggering receptor expressed on myeloid cells 2.

### Processing for electron microscopy

Samples used for electron microscopy (EM) containing the ventral hippocampus were those already processed for immunoperoxidase staining or unstained ones (in which case, they were washed with PBS to remove cryoprotectant). Samples for transmission (T)EM^11^ were post-fixed in 1% osmium tetroxide (EMS, #19190) in phosphate buffer (PB; 100 mM at pH 7.4) and alternatively, for scanning (S)EM^13,15^, sections were post-fixed with a freshly prepared solution containing 3% potassium ferrocyanide (Millipore Sigma, #P9387) in PB combined with an equal volume of aqueous 4% osmium tetroxide for 1 hour. They were afterwards washed with double-distilled water twice and incubated in a freshly prepared and filtered 1 % thiocarbohydrazide (Millipore Sigma, #223220) for 20 minutes^27^. Tissues were rinsed again with double-distilled water and thereafter placed in 2% osmium tetroxide in double-distilled water for 30 minutes. Following this second exposure to osmium tetroxide the tissues were washed in double-distilled water and dehydrated in increasing concentrations of ethanol. They were treated with propylene oxide (Millipore Sigma, #110205) and then impregnated in Durcupan resin (Millipore Sigma, #44610) overnight. After mounting between ACLAR embedding films (EMS, #50425), the resin was polymerized at 55°C for 72 hours. Brain sections from different animals and developmental stages were processed simultaneously in multi-well dishes to ensure the uniformity of the experimental procedure.

### Electron microscopy imaging and analysis

*TEM*. Areas of interest were excised from the embedding films, re-embedded at the tip of resin blocks, and cut at a thickness of 65-80 nm using an ultramicrotome (Leica Ultracut UC7). Ultrathin sections were collected on bare square mesh grids (EMS). Imaging was performed at 80 kV using a FEI Tecnai Spirit G2 TEM. Pictures of dark microglia were captured at magnifications ranging between 440X and 9300X using an ORCA-HR digital camera (10 MP; Hamamatsu). Profiles of neurons, synaptic elements, microglia, and astrocytes, were identified according to well-established criteria^28^. Dark microglia were strictly identified based on a series of ultrastructural features described previously^11^. Briefly, the dark microglia presented condensed cytoplasm and nucleoplasm. They displayed markers of cellular stress, including dilated endoplasmic reticulum and loss of heterochromatin pattern. The dark microglia also displayed hyper-ramification of their processes that extensively interacted with the brain parenchyma, especially synaptic elements around them. They often showed close proximity and direct juxtaposition to blood vessels.

To assess colocalization of dark microglia with various protein markers labeled by immunoperoxidase staining, imaging was conducted at the tissue-resin border where the penetration of antibodies and staining intensity is maximal. The analysis was strictly conducted in tissue areas where intense immunostaining on dark microglia or nearby cells was observed, to rule out the possibility that these cells were not stained due to a limited penetration of the antibodies.

For densitometry analysis, sections containing the ventral hippocampus CA1 *strata radiatum* and *lacunosum-moleculare* were rigorously screened and photographed at magnifications between 4600X and 9300X, corresponding to a total neuropil surface of ∼300,000 μm^2^. The density of dark microglia and typical microglia was determined according to a method described previously^11^. In brief, to calculate the surface area by imaging the section at the lowest magnification possible under the TEM (440X) to determine systematically the total number of grid squares enclosing tissue from both the *strata radiatum* and *lacunosum-moleculare*. The total surface area was calculated at high precision by multiplying the number of grid squares containing the *stratum radiatum* and *stratum lacunosum-moleculare* pooled together by the area of a single grid square. A total of 81 dark microglia and 330 typical microglia across 3-4 animals/group (P5, P10, P15) were included in the analysis. We did not attempt to distinguish resident microglia from bone marrow-derived macrophages and other types of myeloid cells in the brain, but all microglia analyzed were in the parenchyma.

*SEM*. Areas of interest were excised from the embedding films, re-embedded at the tip of resin blocks, and cut at 65-80 nm of thickness using an ultramicrotome (Leica Ultracut UC7 or ARTOS). Ultrathin sections collected onto silicon chip specimen supports (cat 16007 Ted Pella) and affixed to SEM stubs using double-coated carbon conductive tabs. Stubs were imaged using a Zeiss Crossbeam 540 dual beam SEM or Zeiss Crossbeam 350 SEM running ATLAS5 software to create tiled images over the entire block face. The sections were first imaged at 25 nm/pixel resolution to identify the regions of interest. Regions of interest were subsequently reimaged at 5 nm/pixel resolution to confirm the presence of dark microglia and analyze their features. The images were acquired using the ESB and SE2 detectors of the microscopes using a 5 mm working distance, at a voltage of 1.4 kV and current of 1.2 nA. Individual images were stitched together and exported.

*Volumetric focused-ion beam (FIB)-SEM*. A sample block from the *stratum lacunosum-moleculare* of a P10 animal was prepared for analysis by FIB-SEM. The block was trimmed with a razor blade to expose the region of interest (ROI) and then mounted with silver epoxy on a 45° pre-titled SEM stub and coated with a 4 nm layer of platinum to enhance electrical conductivity. Milling of serial sections and imaging of the block face after each *Z*-slice were carried out with the FEI Helios Nanolab 660 DualBeam FIB-SEM equipped with Auto Slice & View G3 version 1.5.3 control software.

Inside the FIB-SEM, the sample blockface was first imaged to determine the orientation of the tissue and identify the ROI. A protective platinum layer 60 μm long, 7 μm wide, and 2 μm thick was deposited on the surface of the ROI (ventral hippocampus *CA1 lacunosum-moleculare*) to provide a flat conductive surface for milling, i.e. orthogonal to the block face of the volume to be serially milled and imaged. Trenches on both sides of the ROI were created to minimize re-deposition of debris. Fiducial marks were milled for aligning ion and electron beams, and were also used to dynamically correct for drift in the x-and y-directions during data collection by applying appropriate SEM beam shifts. Milling was carried out at 30 kV with an ion beam current of 2.5 nA, stage tilt of 4 °, and working distance of 4 mm. At each step, 14 nm of the block face was ablated by the ion beam. Each newly milled block face was imaged with the SEM beam, at an accelerating voltage of 2 kV, beam current of 0.4 nA, stage tilt of 4 °, and working distance of 3 mm. Signal was collected simultaneously with the Through the Lens Detector (TLD) for backscattered electrons and the In-Column Detector (ICD). The x-y pixel size was 13.7 nm with a dwell time of 30 μs per pixel. Pixel dimensions of the images collected were 1536 × 1024, with a total of 396 images were collected.

A dendrite with extensive contacts with the dark microglia was selected from the dataset for visualization. The dark microglia processes, dendrite, and associated axons and synaptic elements were manually segmented and proofread using VAST Lite software. Segmented data was exported as .obj mesh files from VAST Lite and .obj files were imported into Blender. Meshes were processed using BlendGAMer, to provide mesh optimization and smoothing while preserving local geometry. Meshes were further processed using InstantMeshes software, which reduced vertex counts while preserving geometry by producing field-aligned meshes. Meshes were visualized together with the dataset using the Neuromorph plugin for Blender. Neuromorph was also used to generate contact surfaces between dark microglia and dendritic and axonal processes. Figures and supplemental videos were animated and rendered using Blender. Videos were compressed using Handbrake. Post-processing was performed using Adobe Premier Pro.

*H01 Dataset Investigation*. The H01 dataset^17^, a newly released data set containing 1 mm^3^ segmented section of a human anterior temporal cortex isolated from surgical resection from a 45-year-old female donor, was explored through its viewer, Neuroglancer, for dark microglia. The included example was from layer 5 (c3 segmentation IDs: 40881443277 and 85062801060; segmentation of this cell is non-contiguous and thus represents multiple segmentation IDs). The interaction of this dark microglia’s processes (c3 segmentation ID: 85062801060) apposed to an axon from a pyramidal neuron in layer 6 (c3 segmentation ID: 5013063676) was visualized in Neuroglancer. Images presented in the manuscript were obtained from Neuroglancer interface.

### Imaging mass cytometry image acquisition, processing and spatial analysis

A panel of 18 antibodies was selected for *in situ* immunophenotyping of dark microglia (**Table 2**). Panel comprises antibodies that were either commercially available with conjugated metal isotopes or underwent in-house metal conjugation using MaxPar X8 labelling kit following manufacturer′s recommendations (Standard BioTools). Flash-frozen, 6μm mouse sections underwent IMC staining using an adapted protocol from manufacturer (Standard BioTools).

**Table 2.**
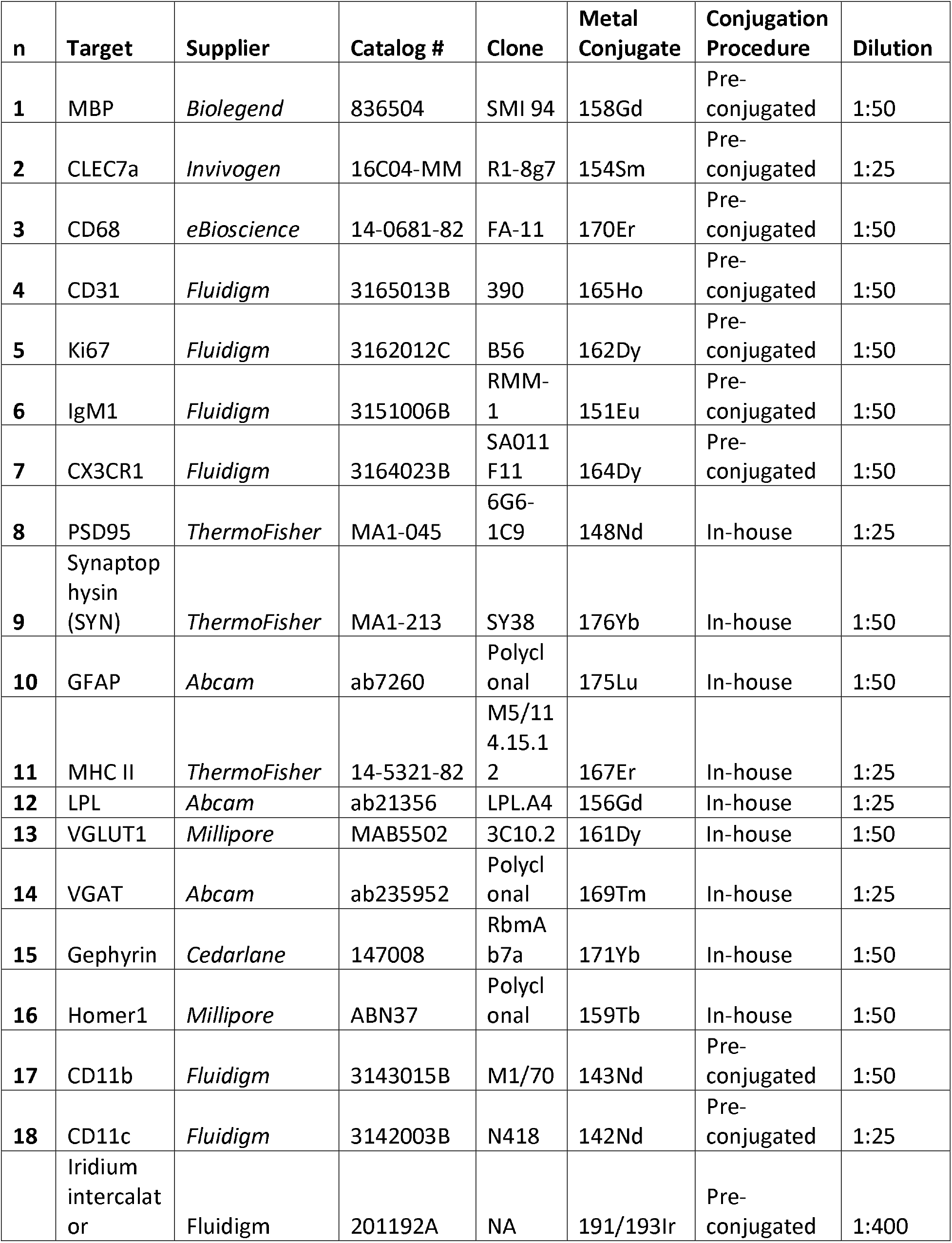
Antibody panel for IMC.

IMC data was acquired on a CYTOF-Helios instrument coupled to a Hyperion Tissue Imager (Fluidigm). Tissue sections including ventral hippocampus CA1 underwent laser ablation at a 1 μm^2^ pixel size/resolution using a frequency of 2001ZHz; resulting files were stored using the MCD format. Raw images were manually inspected before processing using MCD Viewer version 1.0.560.2.

MCD files were processed using a modified version of the Bodenmiller pipeline for multiplexed image analysis ^29^. Briefly, acquisition and channel metadata, as well as corresponding tiff files per each channel within each ROI, were extracted using the *imctools* Python package. Tiffs were then imported onto CellProfiller version 4.0.7 to obtain ilastik-compatible files in the H5 format ^30^. Ilastik version 1.3.3 was used to create probability images for each ROI ^31^. The pixel classifier was trained using manual annotation of DNA iridium-intercalator as nuclear and a combination of glia and immune cell markers (CD11b, CD11c, CD68, CX3CR1 and GFAP) as cytoplasmic signals. Cell masks were then created using CellProfiller ^30^. Nucleated cells were segmented using a 9-20-pixel diameter window as cut-off, whereas cytoplasmic segmentation was achieved by employing an adaptive propagation strategy using the 2-class otsu algorithm.

Corresponding folders containing the combined cell mask, tiff images of markers and background channels were loaded into histoCAT v1.76 ^32^. IMC data was initially arch-sinh transformed in histoCAT. Unsupervised clustering was performed using the Phenograph algorithm based on expression data from the following channels: CD11b, CD11c, CD31, CX3CR1, GFAP, Ki67, Synaptophysin and MBP. Phenograph clusters identified as glia cells were visualized on a t-SNE plot. Heatmap of selected channels and Phenograph-derived clusters, cluster superimposition onto ROIs or Image_id, were also created using histoCAT.

The significance of interactions between Phenograph-derived clusters was assessed using a permutation-based method to analyze single-cell spatial neighborhoods as implemented in histoCAT. Cellular neighborhoods were probed using 999 permutations per analysis and a 10-pixel expansion as a cellular neighbor cut-off.

### Statistics

For quantitative data, except for IMC—which was analyzed as described above, statistics were performed using Prism 10 Software (GraphPad). Data are presented as mean ± standard error of the mean (SEM) and p<0.05 was considered statistically significant. For comparison of dark and typical microglia densities microglia across P5, P10 and P15, a one-way ANOVA was utilized; as there were no overall significant differences there were no *post-hoc* analyses for these comparisons. For comparison between P15 wild-type and TREM2 KO animals, a two-tailed Student’s t-test was utilized. F-or t-vales, p-values and eta-squared (R^2^) are reported in the figure legends.

## Acknowledgements

We acknowledge with respect the lək□⍰⍰ŋ⍰n peoples on whose traditional territory the University of Victoria stands and the Songhees, Esquimalt and WSÁNEĆ peoples whose historical relationships with the land continue to this day.

This work was supported by Canadian Institute of Health Research (CIHR) grants FDN341846 to M-ÉT and PJT461831 to M-ÉT and LP-J. Equipment utilized in this grant was supported by the Canadian Foundation for Innovation (CFI) John R. Evans Leaders Fund and Infrastructure Operating Fund. M-ÉT previously held a Canada Research Chair (Tier II) in Neuroimmunity Plasticity in Health and Therapy at Université Laval and currently holds a Canada Research Chair (Tier II) in Neurobiology of Aging and Cognition at the University of Victoria. UM was supported by NIA P30AG068635 (Nathan Shock Center), Core Grant application NCI CCSG (CA014195), NSF NeuroNex Award (2014862), G. Harold & Leila Y. Mathers Foundation, and the CZI Imaging Scientist Award DOI https://doi.org/10.37921/694870itnyzk from the Chan Zuckerberg Initiative DAF, an advised fund of Silicon Valley Community Foundation (funder DOI 10.13039/100014989). Further support included a National MS Society Research Grant RFA-2203-39318 to LP-J and SP, and a Wellcome Trust Clinical Research Career Development Fellowship G105713 to LP-J. ZSV holds support from the National Institutes of Health (NIH) ROI NS44025.

HAV was the recipient of a CIHR postdoctoral fellowship, a Brain Canada—Canadian Consortium for the Investigation of Cannabinoids (CCIC) Neuroscience Fellowship in Cannabis and Cannabinoid Research, a British Columbia (BC) Women’s Health Research Institute (WHRI) fellowship and was a Michael Smith Health Research BC Research Trainee. K.B. was supported by excellence scholarships from Université Laval and Fondation du CHU de Québec. MEG-S was supported by Medical Research Council Doctoral Training Grant RG86932 and School of Clinical Medicine Cambridge Trust Scholarship salary support. M-KS-P was supported by a doctoral training award from CIHR and M-KS-P, KS and MB were supported by doctoral scholarships from the Fonds de recherche du Québec-Santé (FRQS). J.C.S. was supported by a postdoctoral fellowship from FRQS. KP was also supported by a doctoral scholarship from the Centre de Thématique de Recherche en Neurosciences. CMW received salary support from a National MS Society Post-doctoral fellowship FG-2008-36954. SML, CJM and VL were the recipients of Canadian Graduate Scholarships–Master’s at the University of Victoria. CJM was also the recipient of a graduate grant from the Branch Out Neurological Foundation. T.H. was supported by an Undergraduate Student Research Award from the Natural Sciences and Engineering Research Council of Canada (NSERC).

Funding agencies had no input into the design or execution of this manuscript.

## Author Contributions

HAV, KB, SWN, MET, YYG, ZSV, MJC, UM, LPJ and M-ÉT conceptualized studies. HAV, KB, KS, SWN, MET, MEG-S, M-KS-P, JCS, KP, MB, NV, RG, YYG, JF, SML, CJM, MPK, TH and VL performed experiments. ZSV and MJC contributed reagents. HAV, KB, SWN and MEG-S analyzed data. HAV and MEG-S created figures. HAV and M-ÉT wrote the initial version of the manuscript, with edits from all of the co-authors. M-ÉT, in collaboration with LP-J, carried out funding acquisition, with SP, ZSV, MJC and UM providing additional funding.

HAV, KB, KS, SWN and MET equally contributed to the manuscripts as co-first authors. MEG-S, M-KS-P and JCS equally contributed to the manuscript. M-ÉT is the corresponding author.

---

**Video 1. 3D FIB-SEM example of a dark microglia’s interactions with synaptic elements**.

Video highlighting the ultrastructure and interactions of a dark microglia cell with the surrounding neuropil.

